# Vector manipulation by a semi-persistent plant virus through disease symptom expression

**DOI:** 10.1101/2020.08.19.258178

**Authors:** Diego F. Vasquez, Felipe Borrero-Echeverry, Diego F. Rincon

## Abstract

The greenhouse whitefly (GWF), *Trialeurodes vaporariorum* (Westwood) (Hemiptera: Aleyrodidae) is rarely associated with potato plants yet is the only known vector of the *Potato yellow vein virus* (PYVV). A host shift related with vector’s cognition often requires neural alterations by the virus. However, PYVV, being semi-persistent, is not supposed to directly affect vector physiology. As such, we propose that changes in potato plants caused by PYVV infection should manipulate insect behaviour to increase transmission. Here, we studied the effect of PYVV infection and symptom expression on GWF biological parameters, and attraction towards infected and uninfected potato plants. We compared survival and development rates of GWF nymphs fed with PYVV-infected plants (symptomatic and asymptomatic) and healthy plants under controlled conditions. We also carried out free-choice tests to determine host preference of GWF adults as a function of PYVV infection and disease symptom expression. We found that PYVV infection (both symptomatic and asymptomatic) reduce GWF survival while increasing development time (in symptomatic plants). Combined, a prolonged development time and reduced survival should favour viral uptake and trigger migration of vectors from symptomatic plants short after acquiring the virus. We also found that symptom expression (yellowing) causes significantly greater GWF attraction and establishment compared to healthy or asymptomatic plants. Interestingly, we found that GWF adults that have previously fed on infected plants switch their host preference choosing and establishing on healthy potato plants, which clearly increases horizontal transmission rates. The mechanism through which this behavioural manipulation takes place is not yet well understood. Our results show that symptoms associated with PYVV infection may account for a set of behavioural modifications that make an improbable vector, such as the GWF, into an efficient agent that increases horizontal transmission rates of PYVV.

**Highlights:**

- PYVD reduces the survival of GWF and increases development time when symptoms occur
- PYVD symptom makes potato, a non-host plant, attractive to GWF
- After feeding on infected plants, GWF preference changes to prefer uninfected plants
- PYVV modulates GWF behaviour to enhance horizontal transmission between plants

## Introduction

Viruses rely on hosts for replication, which is always detrimental to the host. As obligate parasites, viruses need to move from one host to another to persist which means that they require efficient mechanisms to enhance transmission rates. Plant viruses face additional difficulties since their hosts are immobile, which reduces the probability of transmission through direct contact between infected and uninfected individuals which is why many plant viruses rely on arthropod vectors (Hamelin, Allen, Prendeville, Hajimorad, & Jeger, 2016; Jia et al., 2018). Vector transmission rates are often positively correlated with within-host multiplication rates and virulence (Pagán, Montes, Milgroom, & García-Arenal, 2014), but virulence may also increase host mortality, reducing the infectious period and transmission rates. In fact, a wide body of evidence supports the trade-off hypothesis, which states that the level of virulence is driven by the optimum balance between within- and among-host parasite fitness traits (Alizon, Hurford, Mideo, & Van Baalen, 2009).

Some vector-borne plant viruses can be transmitted both horizontally and vertically. In such cases, the level of virulence is often negatively correlated with host’s fitness and, thus, expected to vary as a function of the transmission mode: virulence of virus strains selected for horizontal transmission is expected to be higher than that caused by their counterparts adapted for vertical transmission (Lipsitch, Siller, & Nowak, 1996). In the short-term, however, virulence and transmission mode may depend greatly on internal virus-host interactions, which will allow (or impede) viral particles to either multiply repeatedly at a cost to host fitness or reach its reproductive structures relatively harmlessly. Transmission mode is expected to be subjected to disruptive selection, i.e., both extremes vertical and horizontal are benefited against intermediate strains (Messenger, Molineux, & Bull, 1999), which implies that one transmission mode should be triggered according to the level of virulence achieved. While vertical transmission relies mainly on host survival and reproduction, horizontal transmission depends on vector attraction, arrestment and posterior migration to a healthy plant, all of which are, in part, mediated by biotic and abiotic conditions (plant nutrition and defences, climatic conditions, host availability) (Gallet, Michalakis, & Blanc, 2018; Kerry E. Mauck, Chesnais, & Shapiro, 2018; Su et al., 2015).Furthermore, the high level of virulence associated with vector-mediated transmission (i.e., horizontal) should activate mechanisms that increase the probability of vectors carrying viral particles from infected to healthy hosts.

Symptom expression in host plants is usually associated with high levels of virulence and horizontal transmission of vector-borne viruses (Bosque-Pérez & Eigenbrode, 2011; Casteel & Jander, 2013; Kerry E Mauck, De Moraes, & Mescher, 2016). Viruses often manipulate vector behaviour to increase movement between infected and healthy hosts, but the behaviours which need to be manipulated depends on the modality of transmission, which may be persistent, non-persistent, or semi-persistent (Ng & Zhou, 2015). Even though all three benefit by manipulating symptoms to increase vector attraction towards infected plants, they differ in their need to retain the insect vector. Persistent viruses require greater arrestment since insects need to feed for longer periods of time for the virus to reach the gut and allow viral replication inside the vector. Conversely, the non-persistent viruses require low arrestment times since these viruses only persist for a limited time attached to the vector’s mouthparts and need to infect new hosts quickly. Semi-persistent viruses, even though they do not replicate in the vector, may persist for a longer time in the vector’s salivary glands and as such, require specific arrestment times (enough to be acquired, but not degraded). Disease progression may alter many plant processes and traits to achieve these behavioural changes in insect vectors, including changes in coloration, volatile profiles, alter plant defences, and nutritional quality depending on their mode of transmission (Fereres & Moreno, 2009; Hogenhout, Ammar, Whitfield, & Redinbaugh, 2008).

The *Potato yellow vein virus* (PYVV), is the causal agent of potato yellow vein disease (PYVD), a re-emerging epidemic of potato crops in northern South America which reduces yields by up to 50% (Cuadros et al., 2017; Rincon, Vasquez, Rivera-Trujillo, Beltrán, & Borrero-Echeverry, 2019). PYVV infection is transmitted vertically through infected seed tubers and horizontally in a semi-persistent manner by the greenhouse whitefly (GWF) *Trialeurodes vaporariorum* (Westwood) (Hemiptera: Aleyrodidae) (Salazar 2000; Chávez 2009). In South America, potato is normally cultivated above 1500 m.a.s.l. which is not optimal for GWF survival and development (Curry & Pimentel, 1971). Similarly, potato is not a preferred host for GWF under normal conditions, but outbreaks have been observed in association with PYVD (Cuadros et al., 2017). Furthermore, symptom expression of PYVD is associated with reduction in number and weight of tubers produced, while infected, asymptomatic plants vertically transmit the virus without showing such a yield reduction (Guzmán-Barney, Franco-Lara, Rodríguez, Vargas, & Fierro, 2012). Combined, these facts would suggest that GWF should be a poor vector for PYVV and that in order for it to be attracted towards infected potato plants, symptoms associated with PYVV virulence should be strong modulators of GWF behaviour.

We hypothesize that symptom expression of PYVD increases the attraction of GWF towards infected potato plants, but that arrestment should be low in symptomatic plants. Here, we studied the effect of symptom expression on population parameters and behavioural manipulation of GWF. We show that symptom expression has a negative effect on GWF development and survival. We also show that symptom expression differentially modulates GWF behaviour, depending on whether the vector has fed on healthy or infected plants. To our knowledge, this is the first report of a semi-persistent virus manipulating the behaviour of vector insects.

## Materials and methods

### Vegetable material

*Solanum tuberosum* Phureja group cv., (the Colombian Creole potato) plants were used for all experiments. Virus-free plants were obtained from *in vitro* culture to prevent contamination. Mini-tubers were planted in a soil-rice husk substrate (3:1) in 5.5 L pots and plants were grown under greenhouse conditions (22-35°C, 4-6 lux, 50-70%RH) inside a mesh cage (mesh size 1.35 mm) to avoid infestation by insect vectors that could carry PYVV until the fourth week after sowing (code 105 in potato’s BCCH scale). Plants were then transferred to environmental chambers (Sanyo® MLR 351) set to the required conditions (16 °C±1, 4 lux, and 55±10% RH). Plants were watered twice per week until reaching field capacity.

Infected plants came from symptomatic plants (intreveinal yellowing of primary and secondary veins on the upper third of the plant) collected in the municipality of Subachoque (Cundinamarca, Colombia) (4.978093, -74.155993). Symptomatic field-collected plants were taken to the laboratory and used as donor plants to infect virus-free three-week-old plants, according to the procedure established by Vargas (2010). Newly infected plants were subjected to the same conditions in the environmental chambers as virus-free plants (16 °C±1, 4 lux, and 55±10% RH). Both infected and virus-free plants were subsequently evaluated for the presence or absence of the virus by RT-PCR according to the protocol established by Hernandez-Guzmán and Guzmán-Barney (2014). Plants were classified as symptomatic, asymptomatic, and virus-free plants for the experiments.

### Whitefly rearing

GWF adults were obtained from the colony at AGROSAVIA’s entomology lab, (Tibaitatá Research Centre, Mosquera, Colombia). GWF was reared on bean plants (*Phaseolus vulgaris* L.) in order to avoid contact with a possible PYVV host in an isolated room in a glasshouse (22-35°C, 4-6 lux, 50-70%RH). Viruliferous insects were obtained by allowing them to feed on symptomatic plants obtained from the field, confirmed through RT-PCT. Transmission of PYVV by action of viruliferous GWF was by releasing 30 newly emerged GWF adults on leaves of plants expressing PYVD symptoms. Whiteflies were kept in clamp cages on the underside of symptomatic potato leaves for 48 hours. Insects were then transferred to virus-free plants using the same method to infect new plants. Insects could feed for 48 hours and were then removed. To confirm the presence of PYVV in the plants, RT-PCR was performed as described below.

### Life cycle parameter bioassays

In order to determine the effect of PYVD symptoms on development and survival rates of GWF individuals, an experiment with three treatments in a completely randomized design was established. Healthy, infected symptomatic and infected asymptomatic plants were placed in environmental chambers at 16 °C±1, 4 lux, and 55±10% RH. Ten male-female pairs of GWF adults were released in leaf cages on the underside of three randomly-selected leaflets of each plant. Each treatment consisted of eight plants. After 48 hours, adults were removed from the plants and the number of live and dead nymphs, and development stage (eggs, nymphs, adults) were registered daily for 60 days or until all immature GWF emerges as adults or died.

### Free-choice bioassays

Free-choice bioassays were carried out under controlled conditions (16°C±1, 4 Lux, 65%-75% RH) at the Tibaitatá Research Centre. GWF were collected from the colony and kept in a 5 mL cup. GWF were then transferred to a freezer (4°C±1) for five minutes to reduce their mobility. Virus-free, symptomatic and asymptomatic leaves were kept in vials with water inside mesh cages (1 x 1 x 0.8 m) equidistant in a triangular arrangement, 30 cm apart. Ten GWF adults were then released in the centroid of the triangle, and the number of adults on each leaf was evaluated after 30, 60, 120, 240, and 1440 minutes. Each trial was repeated 30 times using different leaves and insects (300 insects total). Bioassays were preformed using both non-viruliferous and viruliferous insects to determine if there were host preference changes due to virus uptake.

### RT-PCR detection

RNA extraction was carried out using the protocol described by Hernandez-Guzmán and Guzmán-Barney (2014). All RNA extracts were obtained using Trizol® (Invitrogen) according to the manufacturer’s instructions. The cDNA synthesis was carried out by mixing 2 µL of 1X reaction buffer, 0.5 µL of 1 mM dNTPs, 1 µL of 10 mM DTT, 0.5 µL of 0.4 μM of 3’ reverse primer, 0.4 μL of RNase inhibitor, 0.5 μL of MMLV HP and 100 ng of RNA for each reaction. The mix was kept at 42 °C for one hour followed by denaturation at 70°C for 10min.

The PCR reactions contained 2 μL of cDNA, 2 µL of 1X NH4 buffer, 1 µL of 25 mM MgCl2, 0.4 µL of 10 μM dNTPs, 0.4 µL of each forward, and reverse primers to obtain a final volume of 10 μL. The amplification program was set to an initial denaturation at 94 °C for 3 min, 35 cycles of denaturation at 94°C for 1 min, alignment at 55°C for 1 min and extension at 72°C for 1 min followed by a final extension at 72°C for 10 min. As a positive control, a leaf sample of potato plants expressing PYVD symptoms was included. As negative controls, a leaf sample of a cape gooseberry plant (*Physalis peruviana*) infected with *Tobacco mosaic virus* (TMV), and a virus-free potato leaf sample (obtained by *in vitro* meristems culture).

### Data analysis

To compare the average development times of nymphs that fed on symptomatic, asymptomatic and virus-free plants, log-logistic models were constructed describing the proportion of emerged adults, as a function of time, *t*:

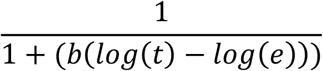

where *b* and *e* are parameters estimated from the data, the former denotes the steepness of the curve and the last equals the midpoint of the s-shaped curve. Parameter *e* (average time at which 50% of adults emerge) was used to compare development times among treatments (healthy, asymptomatic and symptomatic plants). The parameters were estimated by maximum likelihood estimation, assuming a binomial distribution of the response variable. Comparison of estimates of *e* were compared through t-tests.

To assess differences in survival of individuals feeding on the healthy, symptomatic and asymptomatic plants, a generalized linear model with a binomial distribution was fit. The treatment (virus-free, asymptomatic infected and symptomatic infected) was used as an explanatory variable with virus-free plants as the reference factor level. The magnitude of the difference and the significance between the symptomatic, asymptomatic and virus-free levels was examined by Tukey’s post-hoc range test analysis adjusted for generalized models (Bretz, Dette, & Pinheiro, 2010).

For the free-choice bioassays, a multinomial regression analysis was fit for both non-viruliferous and viruliferous GWF to model the proportion of adults selecting each treatment over time. The significance of each model was tested against the null model using a chi-square test. To test for differences within each evaluation time (against the null hypothesis of GWF adults selecting treatments at equal proportions), repeated G-test of goodness-of-fit were performed. G-tests are similar to chi-squared test of goodness-of-fit, but they allow for the inclusion of repetitions (trials), because the G-values of independent repetitions add up to an overall G-value for the whole experiment (Agresti, 2018). Thus, G-values for each trial were estimated and then they were summed for each evaluation time. P-values were calculated from chi-squared distributions using the summed degrees of freedom for each evaluation time. We also performed chi-squared pairwise comparisons of proportions as *poshoc* tests with the data of evaluation times for which significant deviations from equal probabilities were detected with G-tests. Holm’s correction was included to adjust P-values for multiple comparisons (Wright, 1992).

All analyses were carried out using R software (R Core Team, 2020). The package “drc” was used for parameter estimation of development models (Ritz, Baty, Streibig, & Gerhard, 2016), and G-tests for free-choice data were performed using “DescTools” (Signorell et al., 2020).

## Results

### Presence of Potato yellow vein virus (PYVV)

A total of 72 plants were tested. Of the 72 plants, 24 came from in vitro microtubers and all tested negative for PYVV. The remaining 48 plants (24 symptomatic and 24 asymptomatic) came from symptomatic field-collected plants according to the infection protocol above. All 48 plants tested positive for PYVV in RT-PCR analyses. These results confirm that our interpretation of disease symptoms is adequate and that our infection procedure was effective. Of each set of plants, six were used in life parameter assays and the rest were used to obtain leaflets for behavioural experiments.

### Life cycle parameters

Symptom expression had a significant effect on the life cycle of GWF. Nymphs fed on infected plants with yellowing symptoms took longer to develop into adults (58.643 ± 0.147 days) than those fed on healthy (53.086 ± 0.1697 days) or asymptomatic plants (50.571 ± 0.109 days). Interestingly, GWF fed on infected asymptomatic plants developed significantly faster than those fed virus-free plants (Figure 1A).

**Figure 1:**
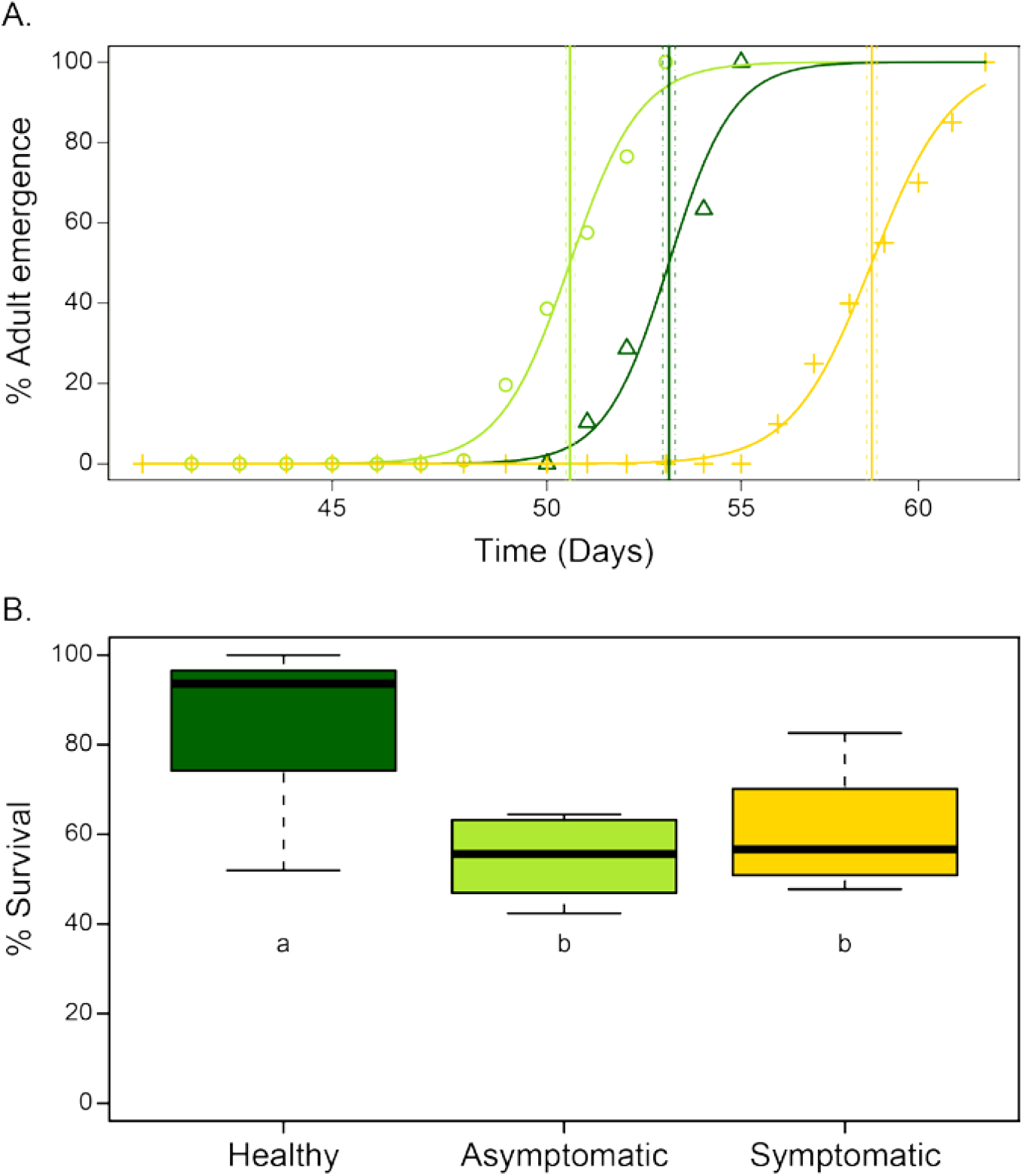
Adult emergence and survival. A. Time in days needed by GWF to reach adulthood when reared on healthy potato plants (dark green), PYVV infected, asymptomatic plants (light green) and PYVV infected, symptomatic plants (yellow). Lines represent the average emergence time and the 95% confidence interval. B. Percent of nymph survival to adulthood when reared on healthy potato plants (dark green), PYVV infected, asymptomatic plants (light green) and PYVV infected, symptomatic plants (yellow).

We found that survival rate of GWF nymphs was significantly affected by PYVV infection (*X*^2^ = 81.059, *P* < 0.001). In fact, the *poshoc* analysis revealed that PYVV infection affected survival of GWF nymphs regardless symptom expression, since survival was equally reduced in nymphs fed with either symptomatic (mean survival = 60.702 ± 4.414 %) or asymptomatic plants (mean survival = 54.803 ± 3.079 %), compared to those that were fed in healthy plants (mean survival = 85.072 ± 6.186 %) (Figure 1B).

### Free-choice bioassays

### Non-viruliferous GWF host choice and settlement

Host-plant preference of GWF adults without previous exposure to PYVV was consistent throughout the observation time, according to the multinomial regression model (*X*^2^ = 2,300, *P* = 0.3166). GWF adults consistently preferred symptomatic, followed by asymptomatic and rarely chose, and settled on healthy leaflets (Figure 2A, Table 1).

**Table 1.** Repeated G-tests of goodness-of-fit for each evaluation time for a free-choice test with adults of *Trialeurodes vaporariorum* with no previous contact with PYVV. The G-value presented is the summed value of independent repetitions for each time. Degrees of freedom (DF) differ among times because the trials with no responses were removed from the analysis (*N* = 30).

|  | T1 (30 min) | T2 (60 min) | T3 (120 min) | T4 (240 min) | T5 (1440 min) |
| --- | --- | --- | --- | --- | --- |
| G-value | 52.524 | 94.833 | 92.319 | 151.339 | 124.515 |
| DF | 32 | 60 | 60 | 60 | 60 |
| *P* | 0.0125 | 0.0027 | 0.0046 | < 0.0001 | < 0.0001 |

**Figure 2.**
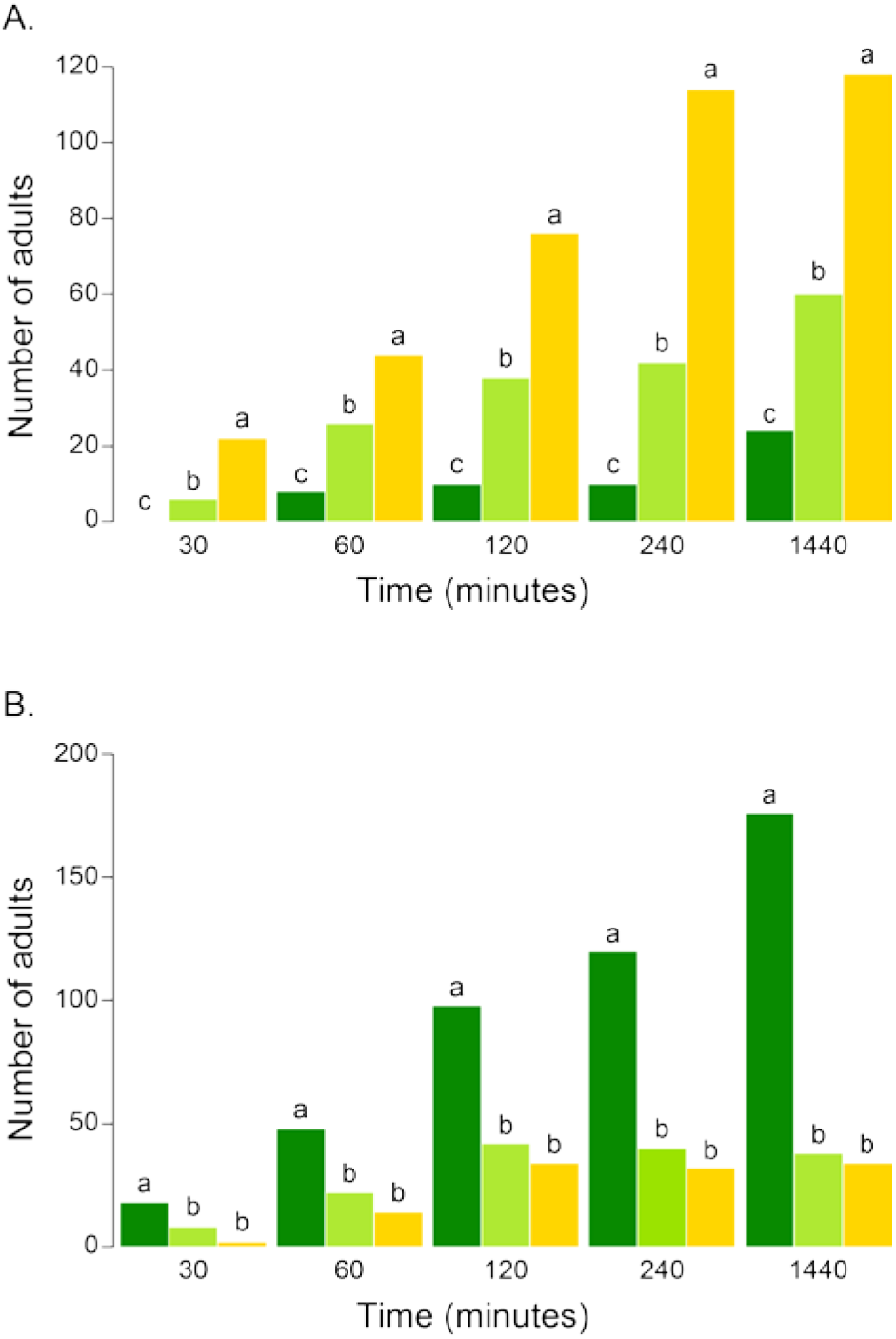
Host plant preference of GWF adults over time. A. Host plant choice of non-viruliferous (previously fed on bean plants) GWF adults in three-way experiments. B. Host plant choice of viruliferous (previously fed on PYVD symptomatic potato plants) GWF adults in three-way experiments. Dark green bars represent healthy potato plants, light green bars represent PYVV infected, asymptomatic plants and yellow bars represent PYVV infected, symptomatic plants. Different letters above the bars denote significant differences, according to chi-squared pairwise comparisons (*P* < 0.05), which were performed for each evaluation time independently.

### Viruliferous GWF host choice and settlement

GWF adults that had previous experience feeding on PYVV-infected plants showed a contrasting host-plant preference in relation with conspecifics without such experience. We found that the proportion of insects that chose healthy leaflets over symptomatic or asymptomatic leaflets increased over time (X^2^ = 8.992, *P* = 0.011) (Figure 2B). No differences were detected in preference for PYVV-infected leaflets, whether they were symptomatic or not (Figure 2B, Table 2).

**Table 2.** Repeated G-tests of goodness-of-fit for each evaluation time for a free-choice test with adults of *Trialeurodes vaporariorum* with previous contact with PYVV. The G-value presented is the summed value of independent repetitions for each time. Degrees of freedom (DF) differ among times because the trials with no responses were removed from the analysis (*N* = 30).

|  | T1 (30 min) | T2 (60 min) | T3 (120 min) | T4 (240 min) | T5 (1440 min) |
| --- | --- | --- | --- | --- | --- |
| G-value | 50.431 | 92.070 | 99.6371 | 146.736 | 215.992 |
| DF | 28 | 56 | 60 | 60 | 60 |
| *P* | 0.0057 | 0.0017 | 0.00099 | < 0.0001 | < 0.0001 |

## Discussion

It is difficult to understand how a generalist insect, such as the GWF, could begin to actively prefer a non-host plant. In order for this to happen, either the insect would need to change its preferences to include the previously non-host plant, or the plant would need to begin giving off the proper cues for the insect to begin finding it attractive. While this kind of cognitive change is less likely to occur and establish itself in an insect population (Libersat, Kaiser, & Emanuel, 2018), it is common for plant’s metabolism to be altered by biotic (diseases, herbivory, phenology) and abiotic (drought, nutrient stress) factors in such a way that the cues they produce change substantially. Such cues (often associated with symptom expression) may include changes in plant coloration and leaf structure (Lu et al., 2017), volatile profile (Fereres et al., 2016), nutritional quality (Bosque-Pérez & Eigenbrode, 2011; Kerry E. Mauck et al., 2018; Szczepaniec & Finke, 2019), and/or nutrient allocation (Byrne & Bellows Jr, 1991; Fereres, 2015; Szczepaniec & Finke, 2019).

Physical and chemical cues generated by virus-infected hosts are fundamental to the behavioural manipulation of insect vectors. Behavioural manipulation of insect vectors through symptom expression has been widely studied in different plant-virus families (Colvin et al., 2006; Fereres et al., 2016; Van Roermund & van Lenteren, 1992) including criniviruses (Jones, 2003; Martelli et al., 2002; Navas-Castillo, López-Moya, & Aranda, 2014; Osorio et al., 2016). Yellowing, the most noticeable symptom of PYVD, plays an important role in turning potato plants from being unattractive for GWF into a potential host. Like many hemipterans, GWF is attracted to yellow (Vaishampayan, Kogan, Waldbauer, & Woolley, 1975) so it could be predicted that yellowing would increase attraction of GWF to a non-plant. Our results show that this is, in fact, the case. Non-viruliferous GFW adults significantly prefer leaves that express PPYD yellowing over green leaves and remain and feed on them in approximately constant proportions, at least over the duration of our experiment (1440 minutes). By doing so, they accomplish the first step required for there to be horizontal transmission of PYVV which is the uptake of the virus. Curiously, leaves from infected, asymptomatic plants show an intermediate attraction and arrestment of GWF adults, suggesting that while not obvious to us, there are more symptoms at play that we are unable to easily detect, such as changes in the volatile profile of plants. It remains to be seen if these unseen symptoms of infected plants that have not begun to express yellowing are enough to attract GWF in the field.

The second key factor for horizontal transmission through vectors to be feasible is arrestment of insects on infected plants in order for them to uptake the virus. According to the postulates of semi-persistent virus transmission, vectors should spend a moderate amount of time feeding on infected plants in order to acquire enough viral titre to establish itself in the insect’s mouth parts and foregut before they move to a new host. The fact that GWF development rate is slower when individuals are reared on symptomatic plants compared to when they develop in healthy plants is likely an effect of reduced nutritional properties of diseased plants, but it may also favour the uptake of viral particles through prolonged feeding duration, likely facilitating horizontal transmission. In fact, such decreased nutritional quality of infected plants may explain why nymphs fed with both symptomatic and asymptomatic plants showed reduced survival compared to those fed with healthy plants (Chesnais et al., 2019). The reduced nutritional quality of infected plants may also be a stimulus for adult GWF to seek out a better-quality host, leading to the third necessary step for horizontal transmission to occur. Curiously, GWF development rate increased when nymphs were reared on infected, asymptomatic plants, suggesting that there may be benefits to feeding on plants that do not express symptoms compared to both healthy and symptomatic plants. Plants being attacked by a virus and an insect, simultaneously, must spend more energy on defences for both. However, if insects arrive on a plant which has already been infected, but whose nutritional quality is not affected yet, it may be able to take advantage of the fact that the plant is already using resources and activating its defences against the virus, and increase its ability to reproduce and develop. Virus-free plants, on the other hand, may concentrate their defences against the insect herbivore, thus reducing their reproductive potential and increasing their development time (Bak, Cheung, Yang, Whitham, & Casteel, 2017; Gallet et al., 2018; Tzanetakis, Martin, & Wintermantel, 2013).

Combined with reduced nutritional quality, we see that viral acquisition also seems to directly affect GWF host preference. Interestingly, although only the proximal parts of the midgut and mouthparts are reached by semi-persistent viruses, PYVV seems to be able to alter vector host preference to increase the dispersion of viral particles from infected to healthy hosts. Similar to what has been observed in other systems such as that of *Tomato severe rugose virus* (ToSRV, Geminiviridae) and *Tomato chlorosis virus* (ToCV, Closteroviridae) (Bosque-Pérez & Eigenbrode, 2011; Casteel et al., 2014; Fereres et al., 2016; Peñaflor, Mauck, Alves, De Moraes, & Mescher, 2016; Wu, Davis, & Eigenbrode, 2014), GWF adults change their host preference as a function of their previous exposure to PYVV-infected plants. Changes in insect host-preference modulated by the pre-acquisition of viruses has been well documented for persistent viruses which replicate inside the vector (He, Li, & Liu, 2015). However, our results show that a semi-persistent virus, through an unknown mechanism, may also modulate insect behaviour in contrast to what has been previously reported in other studies (Whitfield, Falk, & Rotenberg, 2015). This highlights the lack of understanding we have on semi-persistent viruses and how broad the spectrum of characteristics between non-persistent and persistent viruses actually is. It is likely that, while PYVV may not directly affect GWF cognitive behaviour, classical conditioning of GWF adults may explain GWF behavioural manipulation through symptom expression after PYVV infection in potato plants. We hypothesize that the exposure of GWF adults to low-quality hostplants (e.g., infected, symptomatic potato plants) may alter GWF’s further associations of stimuli with preferred hosts. If that is the case, the whole profile of cues associated with PYVV-infected, symptomatic plants will no longer be used to recognize suitable hosts by experienced GWF adults. Effects of previous experience on host-selection by whiteflies, including the GWF, has been documented elsewhere (Lee, Nyrop, & Sanderson, 2010; Shah & Liu, 2013), and classical conditioning has also been reported for other Hemiptera (Stockton, Martini, Patt, & Stelinski, 2016).

In the PYVV-GWF-potato system, symptom expression reduces tuber formation (and vertical transmission rates) (Guzmán-Barney et al., 2012), so it is expected that symptoms are part of an effective strategy to maximize vector-borne (horizontal) transmission. The changes caused by PYVD on host plants are consistent with the hypothesis that the generation of symptoms by viral infection modifies insect vector behaviour and development to enhance horizontal transmission. Our results suggest that semi-persistent viruses have far more complex strategies for horizontal transmission than was previously thought. Even though we have shown that a semi-persistent virus may affect vector attraction, arrestment, and new host plant choice through the expression of symptoms, the mechanisms behind this behavioural modulation remain unclear. Further experiments into symptoms, such as changes in volatile profiles, and their comparisons to those of GWF hosts are crucial to understand the ecology behind the host plant shift observed. Bromatological studies will help us to understand how the virus could manipulate both vector arrestment and release in order to make sure that there is sufficient uptake of viral particles. Lastly, it would be interesting to see what the effects of viral uptake are on GWF physiology and brain chemistry. It remains difficult to understand how a virus which does not persist for long periods of time in an insect, or replicate within, is capable of completely reversing host-choice. Understanding the underlaying mechanism behind this will broaden our understanding of semi-persistent and perhaps break the boundaries of viral classification.

## Conclusions

We evidenced that physiological changes derived from PYVV infection in potato plants alter development, survival and behaviour of the insect vector, the GWF. In particular, the characteristic yellowing seems to be associated with low-nutritional quality (longer immature development times and reduced survival) for the GWF, but quite attractive to GWF adults with no previous exposure to PYVV-infected plants. In contrast, green, healthy plants seem to provide better nutritional quality (shorter immature development times) than infected plants for the GWF and be particularly attractive to adults that had been previously exposed to PYVV-infected plants. Altogether, we present new insights on the ecological relationships between viruses, plants and insect vectors, and how physiological and morphological consequences of viral infections in plants may act as modulators of plant-vector interactions.

## Acknowledgements

We are grateful to Anngie K. Hernandez and Diana M. Torres (AGROSAVIA) who carried PYVV detection tests on potato seed tubers. This study was funded by the Ministerio de Agricultura y Desarrollo Rural de Colombia with government funds allocated to the Corporación Colombiana de Investigación Agropecuaria (AGROSAVIA) (Grant: “Generación y validación de tecnologías sostenibles de producción para incrementar la competitividad de la cadena de la papa en Colombia”). Research support was provided by government funds assigned to AGROSAVIA. The authors assume full responsibility for the interpretation of results and ideas presented in this manuscript.

## Author contribution

All authors contributed equally to the analysis of results, writing and reviewing of the manuscript.

DFV: Contributed to the original idea as well as the hypotheses. Designed, and carried out the experiments. Collected and organized data. Contributed to the statistical analysis.

DFR: Had the original idea and contributed to the consolidation of the hypotheses. Helped in the organization and systematization of the data. Carried out statistical analysis.

FB-E: Contributed to the consolidation of the hypotheses. Contributed to the design of experiments.

## Data Availability

The datasets generated, collected and/or analyzed during the current study are available from the corresponding author on reasonable request, according to institutional guidelines.

